# Impact of Aging on Theta-Phase Gamma-Amplitude Coupling During Learning: A Multivariate Analysis

**DOI:** 10.1101/2025.02.09.637032

**Authors:** Dawid Strzelczyk, Charline Peylo, Nicolas Langer

**Author notes:** CORRESPONDING AUTHOR : Dawid Strzelczyk, Methods of Plasticity Research, Department of Psychology, University of Zurich, Binzmühlestrasse 14, CH-8050, Zürich, Switzerland.

## Abstract

Aging is associated with cognitive decline and memory impairment, but the underlying neural mechanisms remain unclear. Phase-amplitude coupling (PAC) between mid-frontal theta (5 Hz) and occipital gamma (>30 Hz) oscillations is a proposed marker for parallel storage of multiple items in working memory. However, research has mainly focused on young individuals and epilepsy patients, with only a few studies on aging populations. Moreover, these studies have relied on univariate PAC methods, which can be flawed by potential spurious or biased PAC estimates due to non-stationarity of EEG signals. Additionally, these methods typically assess PAC at the level of individual electrodes, potentially overlooking the broader functional significance of theta-gamma coupling in coordinating neural activity across distant brain regions.

To address these gaps, we employed multivariate PAC (mPAC) through generalized eigendecomposition (GED) analysis, which avoids the pitfalls of non-sinusoidal oscillations. 113 young and 117 older healthy participants engaged in a sequence learning paradigm (6423 sequence repetitions, 55’944 stimuli), in which they learned a fixed sequence of visual stimuli over repeated observations, allowing us to track the mPAC during the incremental process of learning.

Behavioral results revealed that younger participants learned significantly faster than older participants. Neurophysiological data showed that mPAC increased over the course of learning in both age groups and could identify fast and slow learners. However, older participants exhibited lower mPAC compared to younger counterparts, which suggest compromised parallel storage of items in working memory in older age. Finally, stratification analysis revealed that mPAC effects persist across performance groups with similar mid-frontal theta levels, suggesting that theta alone does not account for these effects. These findings shed light on the age-related differences in memory formation processes and may guide interventions to enhance memory performance in older adults and slow learners.

## 1. Introduction

The increasing prevalence of physical and cognitive impairments in the aging population presents significant challenges for society (DeCarli, 2003; Grady et al., 2010). Given the association between older age and declines in learning and memory, it is crucial to enhance our understanding of the neural processes underlying successful or impeded learning in older individuals. However, despite mounting evidence of memory deficits in older age, the precise mechanisms contributing to successful or impeded learning remain poorly understood (Cabeza et al., 2018; Ginsberg & Tarantini, 2023; Riddle, 2007). A growing body of evidence highlights the role of neural oscillatory dynamics, particularly theta-gamma phase-amplitude (PAC), in supporting memory processes.

Phase-amplitude coupling between mid-frontal theta (5 Hz) and occipital gamma (>30 Hz) oscillations has been proposed as a neural substrate for working memory, enabling the parallel storage of multiple items by nesting gamma waves into specific phases of slower theta rhythm (Jensen & Lisman, 1996, 1998; J. E. Lisman & Idiart, 1995; J. Lisman & Jensen, 2013; Sauseng et al., 2009, 2010, 2019; Tort et al., 2010). Occipital gamma oscillations have been associated with the encoding of item-specific information in localized cortical assemblies, particularly during visual processing, while theta rhythms integrate information over broader temporal windows, coordinating synaptic plasticity and network-level communication (Axmacher, Henseler, et al., 2010; Heusser et al., 2016; Karlsson et al., 2022). This temporal organization ensures that discrete memory items remain distinguishable while being held simultaneously within a single theta cycle, providing an explanation for the limited working memory capacity of around 3 to 5 items (Cowan, 2001).

Previous empirical studies have shown that stronger PAC has been associated with enhanced learning and memory performance, supporting the binding of item-specific information within neural assemblies. In particular, stronger theta-gamma coupling has been linked to improved learning of item-context associations and higher working memory capacity (Chaieb et al., 2015; Karlsson et al., 2022; Sauseng et al., 2009, 2019; Staudigl & Hanslmayr, 2013; Tort et al., 2009). The theta-gamma coupling appears to adapt dynamically to memory demands. For instance, an increase in memory load slows down theta waves, allowing additional gamma waves to nest within a single theta cycle, thereby extending memory capacity (Axmacher, Henseler, et al., 2010; Kamiński et al., 2011). This suggests that more memory items, each represented by a gamma wave, require a longer theta cycle to be bound together as a multi-item memory trace. Experimental modulation of theta frequency using transcranial alternating current stimulation (tACS) further supports this relationship (Vosskuhl et al., 2015; Wolinski et al., 2018). Slowing theta rhythms enhances working memory capacity by accommodating more gamma cycles, while faster theta rhythms diminish capacity by limiting gamma nesting.

However, research on PAC has predominantly focused on young individuals, with only a few studies on aging populations (Karlsson et al., 2022; Papaioannou et al., 2022; Patterson et al., 2024). Furthermore, existing studies have relied on univariate PAC methods, which are not only susceptible to spurious or biased estimates due to the non-stationarity of EEG signals (Cohen, 2017b, 2022), but also typically assess PAC within single electrodes, potentially overlooking the broader functional role of theta-gamma coupling in coordinating signals across distant brain regions. This limitation is particularly relevant given that theta oscillations are thought to originate in the hippocampus, while gamma oscillations are associated with activity in the occipital cortex (Axmacher, Cohen, et al., 2010; Axmacher, Henseler, et al., 2010; Buzsaki, 2011; Buzsáki & Wang, 2012; Karlsson et al., 2022; Sauseng et al., 2019). As a result, traditional approaches may fail to fully capture the spatially distributed dynamics critical for understanding theta-gamma interactions in memory processes (Papaioannou et al., 2022; Sauseng et al., 2009).

To address these gaps, we employed multivariate PAC (mPAC) through generalized eigendecomposition (GED), a signal-processing method that extracts meaningful patterns from multichannel EEG data by maximizing the signal of interest while minimizing noise, which avoids the pitfalls of non-sinusoidal oscillations (Cohen, 2017b, 2022). 113 young and 117 older healthy participants engaged in a sequence learning paradigm, in which they learned a fixed sequence of visual stimuli over repeated observations, allowing us to track the mPAC during the incremental process of learning. Given the role of mPAC in memory formation, we hypothesized that mPAC would increase over the course of learning as more stimuli are held in working memory. Moreover, we expect fast learners to exhibit higher overall mPAC strength compared to slow learners. Finally, the difficulties of older subjects in holding stimuli in working memory and binding them into coherent memory traces will be reflected by lower mPAC strength and a smaller increase over the course of learning compared to young subjects.

## 2. Methods

### 2.1. Participants

In the present study 113 young (age range 19 - 43 years; mean age, 24.63 ± 4.52 years, 45 male, 88 right-handed) and 117 older (58 - 84 years; mean age, 69.1 ± 5.31 years, 53 male, 95 right-handed) participants were recruited. Table 1 shows basic demographic details for both age groups. All participants were healthy, reported normal or corrected to normal vision and no current neurological or psychiatric diagnosis. The young group consisted of graduate students at the University of Zürich or other universities nearby. The older subjects were recruited through announcements in newsletters and during lectures within the Senior-University of Zürich. As a compensation the participants were given course credit or monetary reward (25 CHF/h). This study was conducted according to the principles expressed in the Declaration of Helsinki. The study was approved by the Institutional Review Board of Canton Zurich (BASEC - Nr. 2017 - 00226). All participants gave their written informed consent before participation.

**Table 1.** Demographics information.

|  | *Young* |  | *Older* |  | *t* | *df* | *p-value* |
| --- | --- | --- | --- | --- | --- | --- | --- |
|  | *Mean* | *SD* | *Mean* | *SD* |  |  |  |
| Gender | 113 (68f) | | 117 (62f) | | $\chi^2=0.43$ | | 0.513 |
| Age (years) | 24.63 | 3.02 | 69.04 | 5.31 | -68.18 | 228 | 1.2e-153*** |
| Years of Education | 15.09 | 3.2 | 15.41 | 3.31 | -0.74 | 226 | 0.459 |
| MMSE | NA |  | 29.04 | 1.01 |  |  |  |

Note. SD = Standard deviation. Df = Degrees of freedom. f = female. MMSE = Mini-mental state examination.
\* $p < 0.05$ . \*\* $p < 0.01$ . \*\*\* $p < 0.001$

### 2.2. Procedure

The data used in this study was recorded in the context of a larger project that aims to quantify age effects on eye movement behavior and EEG recordings of resting-state and other task-based paradigms (Popov et al., 2023; Strzelczyk et al., 2023; Strzelczyk & Langer, 2024; Tröndle & Langer, 2024). The data was collected in two experimental sessions separated by a week. Upon arrival, the older participants performed the Mini-mental State Exam (MMSE) in order to screen for cognitive impairment and dementia (Folstein et al., 1975). All participants accomplished a MMSE score above the threshold of 25.

During the EEG acquisition, the participants were comfortably seated in a sound- and electrically shielded Faraday recording cage. The cage was equipped with a chinrest to minimize head movements and a 24-inch monitor (ASUS ROG, Swift PG248Q, display dimensions 531 x 299 mm, resolution 800 x 600 pixels resulting in a display: 400 x 298.9 mm, vertical refresh rate of 100 Hz) on which the experiment was presented. The distance between the chinrest and the monitor was 68 cm.

### 2.3. Sequence Learning Task

A visual sequence learning paradigm initially developed by (Moisello et al., 2013) and considered an important tool in assessing reliable indices of memory formation and learning progress (Steinemann et al., 2016; Strzelczyk et al., 2023; Strzelczyk & Langer, 2024) was used. Through the employment of simplified stimuli, this paradigm has the advantage of mitigating confounding effects of stimuli properties such as memorability, semantic content, and sensory characteristics that typically influence learning efficacy. By enabling within-sequence comparisons among simple standardized stimuli, this paradigm enables tracking the progress of gradual memory formation through both neurophysiological and behavioral markers (Gobet et al., 2001; Langer et al., 2017). The participants were asked to learn a fixed sequence of eight visual stimulus positions (Figure 1). The stimuli consisted of filled white circles (visual angle of 0.84°) and were presented on a computer screen with a bright gray background, positioned equidistant around a ring of fixed eccentricity (visual angle of the distance between center of the screen and the stimulus of 4.21°). Each stimulus was presented for 600 ms with an interstimulus interval of 1300 ms. Before the main task recording, a training task was administered, consisting of 4 stimuli placed on the same 8 locations, in order to familiarize the participants with the tasks and to ensure task comprehension. The participants performed the training task until they correctly recalled all 4 locations. Feedback was provided only during the training task.

**Figure 1:**
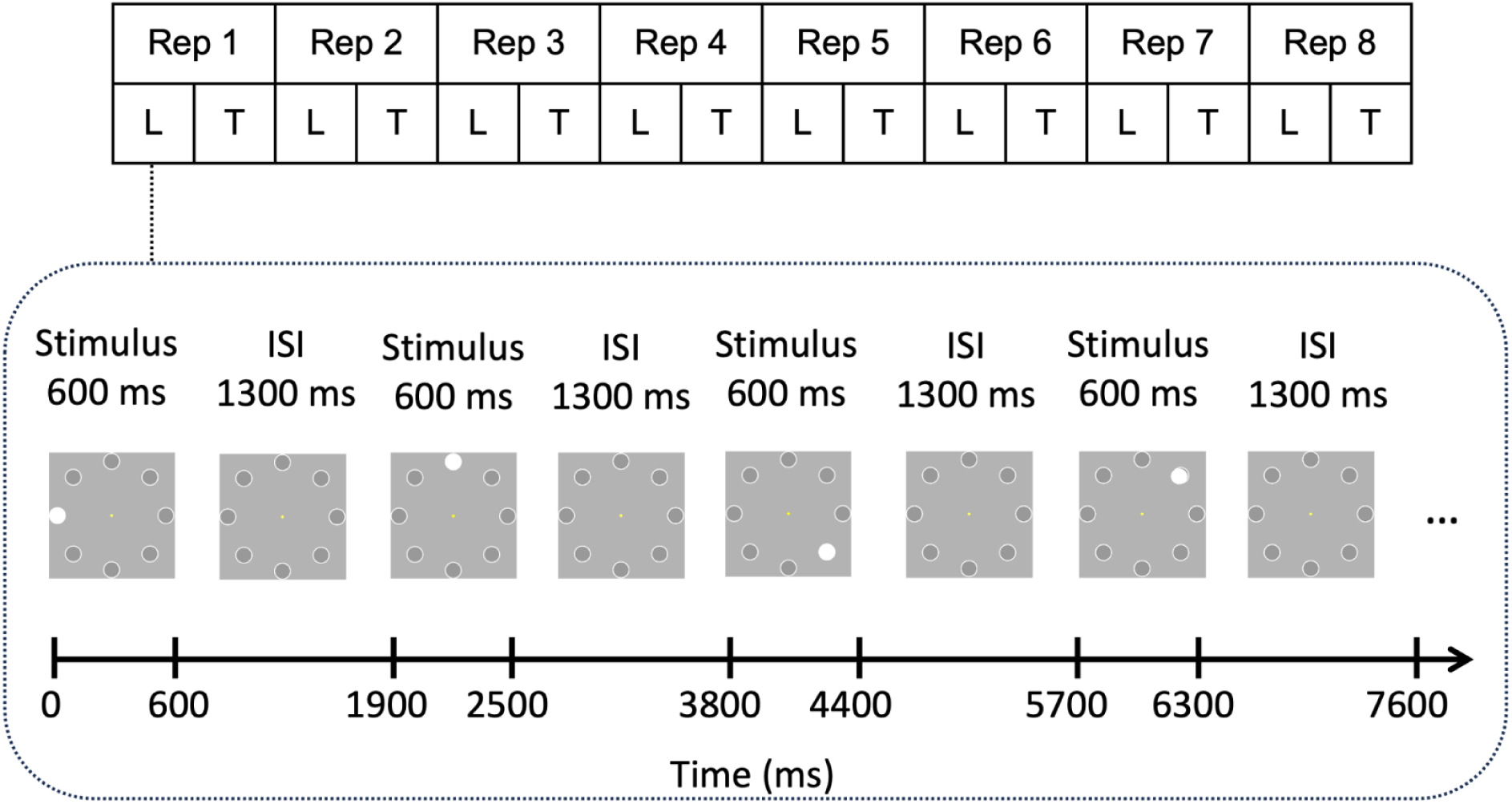
Sequence learning task and the design of the present study. A sequence consisted of eight positions at which white circles were presented one by one. Each stimulus was presented for 600 ms with the interstimulus interval of 1300 ms. A stimulus was presented at each placeholder position during the sequence. Each sequence repetition (Rep) consisted of a learning phase (L) in which all sequence elements were presented and a testing phase (T) in which the participants were asked to recall the position of the stimuli. Adapted from Strzelczyk et al. (2024) licensed under CC BY 4.0.

The main task consisted either of eight sequence repetitions or ended after the participant correctly recalled the sequence of stimuli three times in total (i.e., not necessarily consecutively). Each sequence repetition consisted of a learning phase and a testing phase. In the learning phase, the participants were told to focus on a yellow dot at the center of the screen (controlled by an eye-tracking device) and memorize the position of each stimulus. In the test phase, the participants attempted to recall the sequence using a computer mouse by clicking the locations on a computer screen. There was no time restriction on providing responses, and no feedback on their performance was given. Therefore, the duration for learning a sequence varied between 2 and 5 min, depending on the speed of recall reports and number of sequence repetitions (i.e., 3 to 8 repetitions). Overall, each participant learned 6 different sequences (3 in each session), resulting in a minimum of 18 (in case the participant solves everything correctly from the beginning) and maximum of 48 sequence repetitions in total.

### 2.4. Behavioral Data

To quantify learning performance over sequence repetitions, *accuracy* was computed for each participant. The *accuracy*, which reflects the cumulative sequence knowledge, was defined as the ratio of the number of correct nCorrect(P, sr) responses to the total number of stimuli nT(P, sr) in each sequence repetition, where *P* represents the participant and *sr* the sequence repetition.

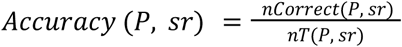

The participants were not instructed to prioritize speed in their responses. Consequently, the reaction time data should be interpreted with caution. For the sake of completeness, however, reaction time analysis can be found in supplementary materials (Supplementary Figure 1).

### 2.5.1. Eye-Tracking data acquisition

An infrared video-based eye tracker (EyeLink 1000 Plus, SR Research; http://www.sr-research.com/) recorded eye movements at a sampling rate of 500 Hz and an instrumental spatial resolution of 0.01°. The eye tracker was calibrated and validated before each task with a 9-point grid until the average error for all 9 points was below 1°. In this study eye tracker data were used in a control analysis accounting for whether the participants maintained focus at the center of the screen as instructed (i.e., yellow dot). Therefore, eye movements during each stimulus presentation period were included into the models as a covariate (i.e., 1 = kept fixation, 0 = lost fixation). It was deemed that a participant had kept fixation, if the gaze was directed at the center of the screen within a square of side 1.26° of visual angle during at least 90% of the total duration of the stimulus presentation period.

### 2. 5. 2. EEG data acquisition and preprocessing

EEG data were recorded at a sampling rate of 500 Hz using a 128-channel Hydrogel net system (Electrical Geodesics Inc.). The recording reference was at Cz (vertex of the head), and impedances were kept below 40 kΩ. All data were analyzed using Matlab 2023b (The MathWorks, Inc., Natick, Massachusetts, United States). Due to technical difficulties (EEG data not saved properly, missing subjects’s responses, Matlab crash during experiment, subject attending only 1 session (i.e., 18 subjects)) data from 118 sequences was not available, resulting in a total of 1262 sequences (i.e., 6423 sequence repetitions, 55’944 stimuli). The data were preprocessed in Automagic 2.6.1, a MATLAB based toolbox for automated, reliable and objective preprocessing of EEG-datasets (Pedroni et al., 2019). In the first step in Automagic, the bad channels were detected using the PREP pipeline (Bigdely-Shamlo et al., 2015). A channel was defined as bad based on 1) extreme amplitudes (z-score cutoff for robust channel deviation of more than 5), 2) lack of correlation (at least 0.4) with other channels with a window size of 1s to compute the correlation, 3) lack of predictability by other channels (channel is bad if the prediction falls below the absolute correlation of 0.75 in a fraction of 0.4 windows of a duration of 5s), 4) unusual high frequency noise using a z-score cutoff for SNR of 5. These channels were removed from the original EEG data. The data was filtered using a high-pass filter with 0.5 Hz cutoff using the EEGLAB function pop_eegfiltnew (Widmann & Schröger, 2012). Line noise was removed using a ZapLine method with a passband edge of 50 Hz (de Cheveigné, 2020), removing 7 power line components. Next, independent component analysis (ICA) was performed. However, as the ICA is biased towards high amplitude and low frequency noise (i.e., sweating), the data was temporarily filtered with a high-pass filter of 1 Hz in order to improve the ICA decomposition. Additionally, we used optimized ICA training data (OPTICAT) for ocular correction, enhancing ICA’s ability to detect and isolate ocular artifacts (Dimigen, 2020). Using the pre-trained classifier IClabel (Pion-Tonachini et al., 2019) each independent component with a probability rating >0.8 of being an artifact such as muscle activity, heart artifacts, eye activity, line noise and channel noise were removed from the data. The remaining components were back-projected on the original 0.5 Hz high-pass filtered data. In the next step, the channels identified as bad were interpolated using the spherical interpolation method. Finally, the quality of the data was automatically and objectively assessed in Automagic, thus increasing research reproducibility by having objective measures for data quality. Using a selection of 4 quality measures the data was classified into three categories: Good (1091 sequences), OK (146 sequences) or Bad (25 sequences). Data was classified as bad, if 1) the proportion of high-amplitude data points (>30μV) in the signal is greater than 0.3, or 2) more than 30% of time points show a variance greater than 30μV across all channels, or 3) 30% of the channels show variance greater than 30μV, or 4) the ratio of bad channels is greater than 0.3. For the further analysis only the datasets with Good and OK ratings were used.

Subsequently, 23 channels were excluded from further analysis, including 10 EOG channels and 13 channels located on the chin and neck as they capture little brain activity and are mostly contaminated with muscle artifacts (Langer et al., 2017; Tröndle et al., 2022, 2023; Tröndle & Langer, 2024). Next, the data was re-referenced to average reference and segmented from −750 to 1150 ms after stimulus onset (i.e., presentation of white circle on one of the eight positions). The segments were inspected using an amplitude threshold of 90μV resulting in 10.7 % rejected segments on average for each participant.

### 2. 5. 3. Multivariate mid-frontal theta-gamma phase-amplitude coupling

We proceeded with the computation of multivariate cross-frequency phase-amplitude coupling using general eigendecomposition as outlined by (Cohen, 2017b, 2022). The procedure is summarized in Figure 2. First, we isolated mid-frontal theta using spatial filters optimized to maximize power within the theta band. This involved finding channel weights that best differentiate theta-band activity from broadband multichannel activity, measured using covariance matrices (de Cheveigné & Arzounian, 2015; Duprez et al., 2020; Nikulin et al., 2011). This process can be described using the Rayleigh quotient, which aims to find a set of channel weights in vector *w* that maximizes:

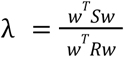

**Figure 2:**
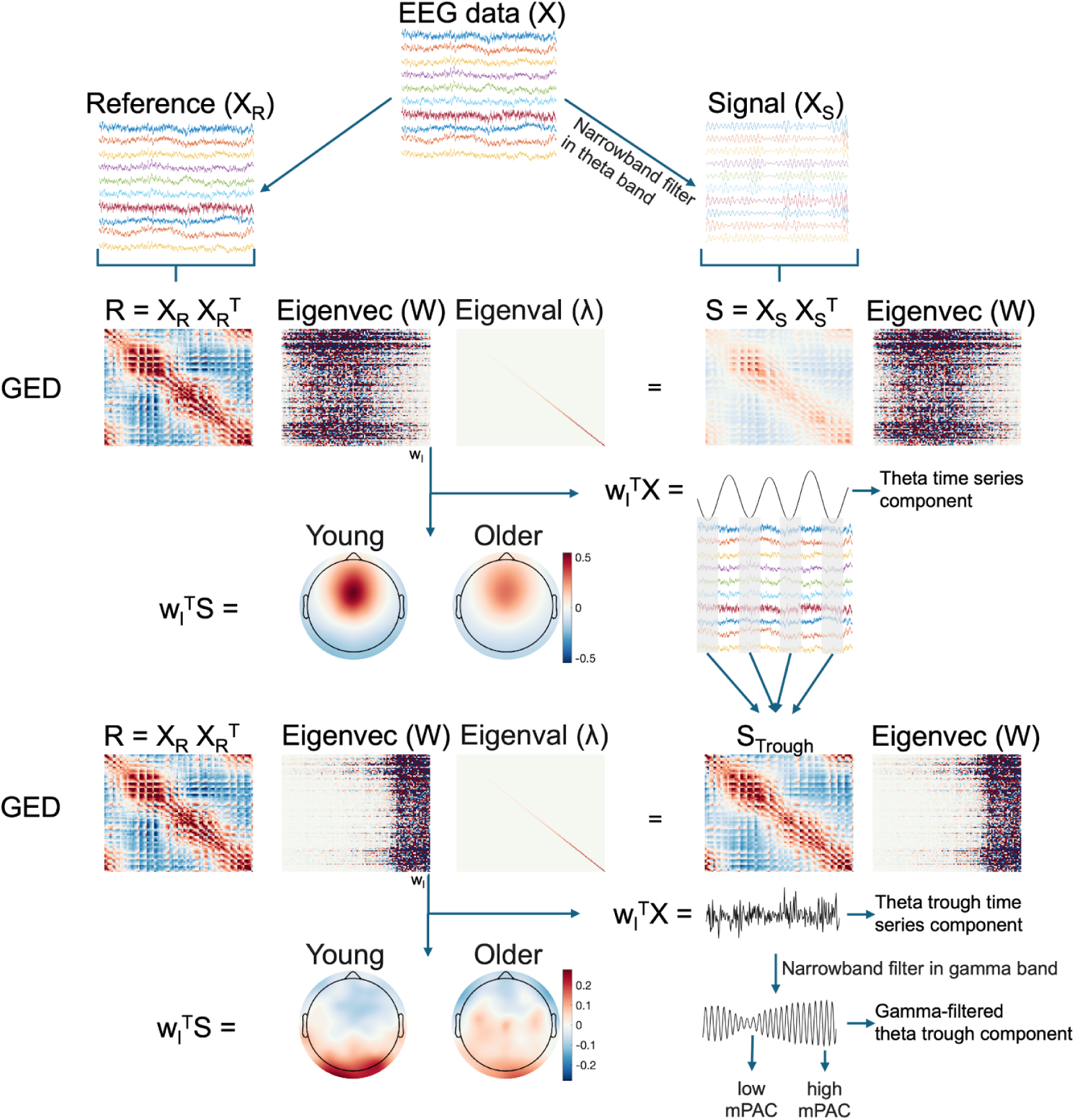
Multivariate theta-gamma phase-amplitude coupling via general eigendecomposition. From EEG data the reference (X_R_) and signal (X_S_) were selected. The X_R_ was defined as the broadband data, while Xs was narrowband filtered in theta band. GED was performed on the two covariance matrices created from the X_R_ and X_S_. The resulting eigenvectors (W) are spatial filters, while the corresponding eigenvalues indicate the ratio of matrix S to R along each w_i_. The eigenvector with highest eigenvalue, when multiplied by data (X), produces a theta time series component and when multiplied by signal covariance matrix, the spatial distribution of the component. Next, a covariance matrix, S_trough_, was formed using broadband data from a 1/4 of a theta-frequency cycle surrounding each theta trough. The R covariance matrix was formed using the entire broadband time series. GED was performed to maximize the difference between broadband signal in theta troughs and the broadband signal. The eigenvector in matrix W with the largest eigenvalue represents the spatial filter that can be used to obtain the theta trough component. This component is a weighted sum of all electrodes that maximally differentiates activity during theta troughs from activity during all theta phases. Finally, the theta trough component was filtered in the gamma range (30–100 Hz, in 2 Hz steps), and mPAC was computed as the difference between peaks and troughs of the gamma-filtered theta trough component at each frequency step within a predefined 500 ms sliding window applied across the trial length (similar to simply computing envelope of the gamma-filtered theta trough component).

In this context, *S* (i.e., signal) represents the covariance matrix obtained from the theta narrowband-filtered data, while *R* (i.e., reference) represents the covariance matrix from the broadband signal. The solution is achieved through a generalized eigenvalue decomposition of these two covariance matrices, expressed by the equation 𝑆𝑊 = 𝑅𝑊λ. This technique is recommended to enhance power at low frequencies (Cohen, 2017a; Duprez et al., 2020; Nikulin et al., 2011) and offers several benefits: it improves the signal-to-noise ratio, accounts for inter-individual topographical differences, and eliminates bias related to electrode selection. The column of *W* corresponding to the highest generalized eigenvalue is the spatial filter that maximizes the difference between *S* and R. This spatial filter is then applied to the data as *y* = *w^T^X* where *X* is the data matrix (channels-by-time), and *w* is the selected column of *W*.

The data for the S and R matrices consists of concatenated, artifact-free trials of one stimulus sequence, ranging from 24 to 64 trials depending on the participant’s learning speed. Before computing the theta and broadband covariance matrices, the data was mean centered (Cohen, 2017b, 2022) to prevent the GED solutions from being biased towards these offsets rather than the direction that optimizes the desired criterion. Mean-centering involves ensuring that the average value of each channel, over the time window used to compute the covariance matrix, is zero. To define the peak frequency of the temporal filter for the S covariance matrix, we determined the subject-wise frequency of peak power in the theta band at electrode FCz, within the range of 4 to 8 Hz to account for variability in theta peak frequency (Cohen, 2014; Duprez et al., 2020). This peak frequency was used as the center frequency for a Gaussian narrow-band filter (FWHM = 4 Hz) applied to the broadband data. The spatial filters’ activation patterns were inspected via topographical maps, with a focus on mid-frontal theta centered around FCz (Duprez et al., 2020; Haufe et al., 2014; Klimesch et al., 2005; Sauseng et al., 2010). To identify the GED component reflecting mid-frontal theta, we simulated mid-frontal theta as a Gaussian distribution centered around FCz with a variance of 4 and computed spatial correlations with the first five GED components, selecting the component with the highest correlation. The theta time series component was derived by multiplying narrowband-filtered theta band data by the eigenvector with the largest eigenvalue, representing the best theta component obtained via spatial correlation (Cohen, 2022).

Having computed the theta time series component, we aimed to identify the brain activity patterns associated with mid-frontal theta troughs (i.e., theta trough network) according to Cohen (2017b). The core idea behind this analysis is based on the similarity between the theta trough and broadband covariance matrices. What is unique, according to previous research, is likely the gamma amplitude observed in peaks and troughs (Cohen, 2017b, 2022). GED maximizes the difference between theta trough and broadband (which corresponds to theta peak), thereby highlighting the gamma modulation between theta peaks and troughs. Here, the S covariance matrix was constructed using broadband data from 1/4 of a subject-specific theta-frequency cycle around each theta trough (1/8 of a cycle before and 1/8 after the trough), while the R covariance matrix was formed using the entire broadband time series. Importantly, the theta frequency was individualized for each participant (as detailed above), meaning that the duration of 1/4 of a theta cycle varied between individuals, depending on their specific theta frequencies. The eigenvector in matrix W with the largest eigenvalue (diagonal elements of matrix ⼊) serves as the spatial filter for extracting the theta trough component. This component is a weighted sum of all electrodes that maximally differentiates the activity during theta troughs from the activity during all theta phases (Cohen, 2017b, 2022).

Finally, we quantified the multivariate phase-amplitude coupling strength. We applied narrowband temporal filters to the theta trough component to compute the average peak-to-trough amplitude differences across frequencies between 30 and 100 Hz in 2 Hz steps. This frequency range was selected to eliminate both theta activity and higher-frequency activity (i.e. >100Hz) unlikely to reflect brain dynamics. The filter’s FWHM values of 5 Hz and 10 Hz produced similar results (Supplementary Material 2); we report the 10 Hz results in the main paper. Next, we segmented the component to the original epochs, ranging from −750 to 1150 ms post-stimulus. For each filtering step, we used a sliding window of 500 ms with a 50 ms step to calculate the mPAC strength by determining the amplitude difference between peaks and troughs of the gamma-filtered theta trough component. This approach provided mPAC strength estimates between −500 and 900 ms, and across the frequency range of 30 to 100 Hz. As a control, we varied the sliding window length for PAC computation, testing windows of 200, 300 and 500 ms to assess the robustness of our findings across different temporal resolutions (see Supplementary Material 3). The chosen time windows ensured that the shortest duration (200 ms) captured at least five full gamma cycles, while the longest window (500 ms) provided sufficient temporal precision without excessive smearing of mPAC estimates over time. The results across all time windows remained highly consistent, therefore we reported the results with a sliding window of 500 ms in the main manuscript.

## 2. 6. Statistical analyses

We analyzed the behavioral and neurophysiological data using linear mixed effect models. In the following formulas fixed effects are denoted by a “+” symbol and interaction effects by an “∗” symbol, in line with the Wilkinson notation (Wilkinson & Rogers, 1973). The predictors included the repetition number (continuous variable: 1-8), age group (factor of 2 levels: young, older), performance group (continuous variable: 3-8), session (factor of 2 levels: 1, 2), eye movements (factor of 2 levels: kept, lost fixation), subject (factor), sequence number (factor of 6 levels: 1-6, i.e., each participant attempted to memorize six sequences).

### 2. 6. 1. Behavioral analysis

We began the behavioral analysis by examining the learning progress of both age groups. Learning performance was formally modeled as the cumulative knowledge about the sequence (i.e., accuracy). We specified a linear mixed effect model with an accuracy as dependent variable (continuous variable: 0-1). The fixed effects included the sequence repetition number, age group, interaction of sequence repetition number and age group, session, ET and the random effects included subject and sequence number.

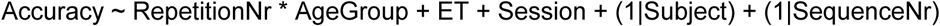

### 2. 6. 2. EEG data analysis

We obtained mPAC values across a frequency range of 30 - 100 Hz and a time window spanning −500 ms to 900 ms relative to stimulus onset. mPAC was computed as the difference between peaks and troughs of the gamma-filtered theta trough component at each frequency step, using a 500 ms sliding window across the trial length (similar computing envelope of the gamma-filtered theta trough component). To address the multiple comparisons problem and account for temporal-frequency dependencies in mPAC, we applied an adapted version of the lmeEEG toolbox (Visalli et al., 2024), which enables computationally feasible mixed-model analyses within a mass univariate framework. Our adaptation of the lmeEEG method involved three main steps. First, we ran linear mixed-effects models at each time-frequency point, accounting for crossed random effects. Each LMM incorporated fixed effects for age group, repetition number, and performance group, as well as their interactions, and a random intercept of the subject.

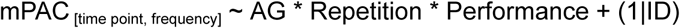

In the second step, we calculated “marginal” EEG by removing random-effects contributions from the data, achieved by combining the fitted values with the residuals. This marginal EEG data allowed each time-frequency point to be analyzed through mass univariate linear regressions, resulting in a map of t-values for each fixed effect. These regressions provided t-values, regression coefficients, and standard errors for each predictor, generating a t-value map at each time-frequency point.

The final step involved assessing significance using cluster-based permutation tests (CBPT). For each permutation, we shuffled mPAC values along three dimensions (frequency, time, and trials) to create a randomized dataset for comparison against the observed data. Next, the test statistic was computed as the maximum of the cluster-level summed t-values, forming an empirical distribution. The observed test statistic was then compared to this empirical distribution to determine the proportion of permutations with a larger test statistic than the observed one, yielding a p-value. If this p-value was below the critical alpha-level (0.05), we rejected the null hypothesis, concluding that significant differences in mPAC existed between experimental conditions. This approach effectively corrected for multiple comparisons while accounting for dependencies across frequency and time. For a more detailed description please refer to (Visalli et al., 2024).

It is important to note that a significant outcome from this test does not identify a specific “significant cluster.” Rather, the test indicates that a statistically meaningful difference exists without specifying precise timing, frequency, or duration of the effect. Thus, identified clusters highlight areas that contributed to rejecting H0, but do not establish exact temporal or frequency boundaries for the observed differences (Maris & Oostenveld, 2007).

### 2. 6. 3. Stratification analysis: matching subjects with similar mid-frontal theta and gamma levels

To ensure that our mPAC effects were not driven by variations in mid-frontal theta or gamma power, we conducted two stratification analyses based on 1) mid-frontal theta and 2) gamma power levels. Stratification analysis is a statistical method used to control for potential confounding variables by dividing a dataset into distinct subgroups, or strata, based on specific characteristics. Within each stratum, the confounding variable is held constant, allowing for a clearer examination of the relationship between the primary independent and dependent variables (Rothman, 2011; VanderWeele, 2009). In this study, we employed stratification to first match mid-frontal theta and then gamma power across different performance groups, thereby controlling for its potential confounding effect on mPAC. Multivariate gamma power was computed similarly to mid-frontal theta power but with a center frequency of 55 Hz and a FWHM of 25 Hz (see Methods 2.5.3). Specifically, we first selected the three performance groups with the highest amount of data (stimulus sequences). As increasing the number of performance groups reduces the pool of subjects with matching power levels, we focused on only the three performance groups with the largest sample sizes (i.e., number of stimulus sequences in a given performance group). We then divided the range of power within these groups into 20 bins between its minimum and maximum values. Within each bin, we randomly selected an equal number of stimulus sequences from the three performance groups, allowing for comparison of mPAC across performance groups among subjects with comparable mid-frontal theta and gamma power. This procedure was repeated 100 times to account for random allocation of stimulus sequences for each mid-frontal theta and gamma bin.

The analysis was conducted separately for young and older adults: young subjects in performance groups 3 (153 sequences), 4 (248 sequences), and 5 (114 sequences), and older subjects in performance groups 5 (106 sequences), 6 (95 sequences), and 8 (223 sequences). Average mid-frontal theta and gamma power used for the stratification analysis was computed as the mean power between 0 and 900 ms across the first three sequence repetitions. Finally, we examined whether differences in mPAC were still evident across performance groups when mid-frontal theta power was matched.

## 3. Results

### 3.1. Behavioral results

Table 1 shows basic demographic information for young and older participants.

#### Visual sequence learning task performance

Across both sessions (i.e., 6 sequences), young subjects required on average 4.45 ± 0.97 repetitions, while older subjects required 6.43 ± 1.10 repetitions to finish the task (M ± SD). The difference was statistically significant (t = −14.27; p = 2.94e-33; CI = [−2.25, −1.70]). The task was considered as finished after 8 repetitions or after the subjects correctly recalled all sequence elements 3 times in total (i.e., fastest learners completed the task after 3 and slowest learners after 8 repetitions).

To investigate learning effects across sequence repetitions, we tested the hypothesis that the knowledge about sequence elements (i.e., accuracy) increases with each sequence repetition and tested for potential age differences. Figure 4 (A & B) shows the progression of accuracy over the sequence repetition. To visualize the interindividual variance in performance, we further subdivided the participants and their sequences into groups based on the number of sequence repetitions required for memorization of all sequence elements (i.e., performance groups). The lines represent the learning curves of each of those groups.

The model for accuracy included fixed effects of repetition number, age group, and their interaction, main effect of eye tracker, and a random effect of subject.

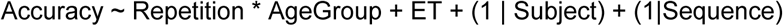

The model revealed a significant main effect of repetition number (β = 0.08; p = 2.4e-94; CI = [0.07, 0.09]), that is the accuracy increased with each sequence repetition (Figure 2A-B). Moreover, there was a significant main effect of age group (β = −0.20 ; p = 1e-18; CI = [−0.25, −0.16]), suggesting a lower accuracy in the older group. There was also a significant interaction of repetition number and age group, indicating a smaller increase in accuracy with each sequence repetition in the older group compared to the increase in the young group (β = −0.01; p = 0.001; CI = [−0.02, −0.01]). Finally, there was a significant main effect of eye movements (β = 0.003; p = 0.023; CI = [0.00, 0.01]), that is the accuracy was slightly higher when subjects were fixating the center of the screen. In addition, we observed a substantial variation of accuracy between subjects (SD = 0.12) and sequences (SD = 0.01). Summarized, young and older subjects gradually learn the stimuli over repeated sequence presentations, with the young learning on average faster than older participants.

### 3.2. Multivariate theta-gamma phase-amplitude coupling (mPAC) over the course of learning

In the next step, we tested the hypothesis that learning progress is associated with changes in mPAC and examined how age influences mPAC. First, we ran mass linear mixed-effects models with crossed random effects for each time point and frequency, incorporating fixed effects of age group, repetition number, performance group, and their interactions, as well as a random effect for individual subjects. Next, we applied cluster-based permutation tests to correct for multiple comparisons, accounting for mPAC dependencies across time and frequency. The results are presented in Figure 3.

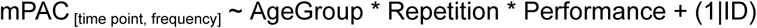

**Figure 3:**
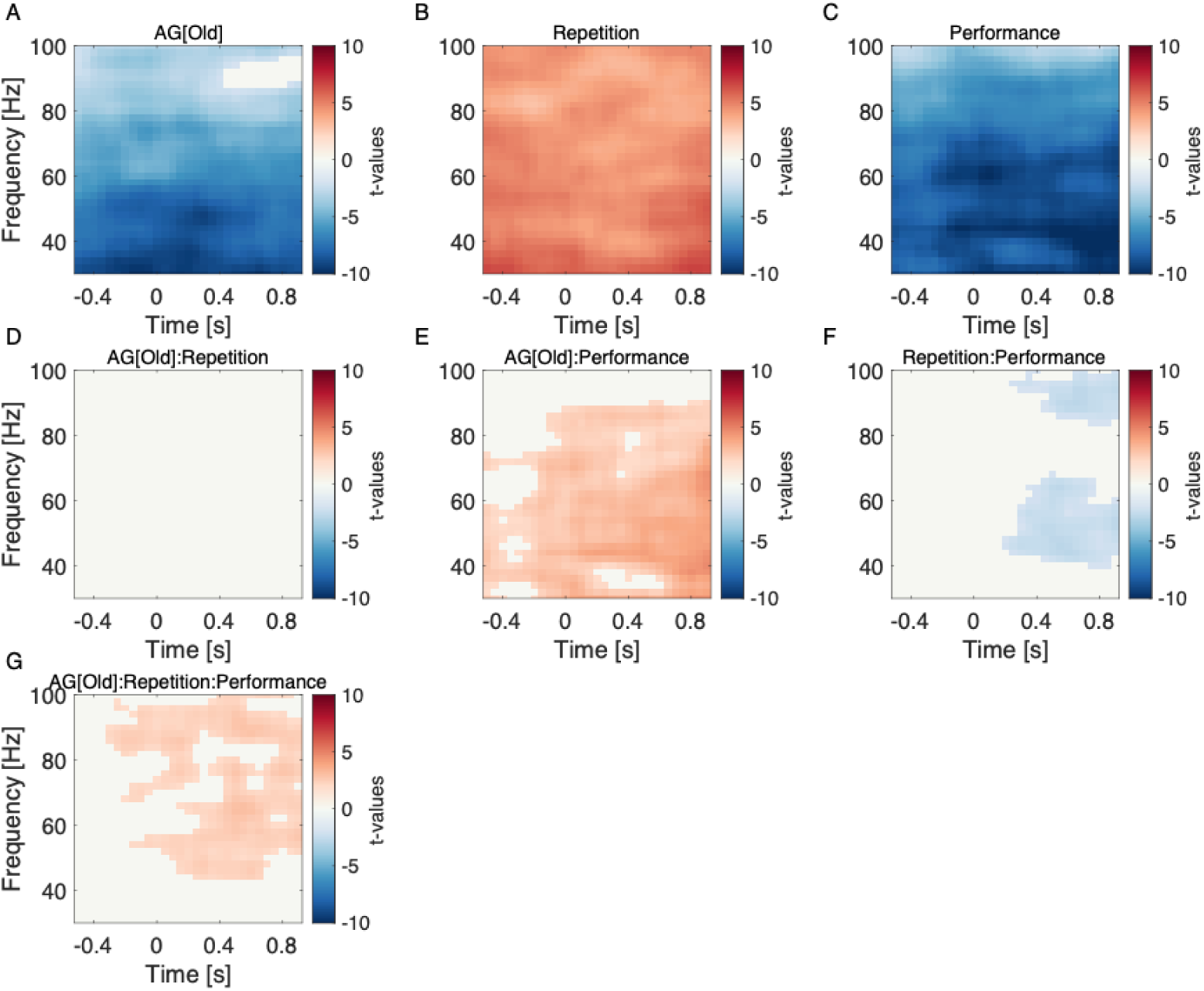
Results of mass linear mixed-effects modeling of multivariate phase-amplitude coupling with crossed random effects. Cluster-based permutation analysis indicated that (A) older adults exhibited lower mPAC compared to younger adults, (B) mPAC values increased across sequence repetitions, (C) slower learners showed lower mPAC compared to faster learners, (E) a significant interaction between age group and performance indicated a stronger relationship between higher mPAC and better performance in younger adults, (F) faster learners demonstrated greater increases in mPAC with repetition, (G) a three-way interaction of age group, repetition, and performance revealed that younger, faster learners had the most pronounced increases in mPAC. Only statistically significant clusters, corrected for multiple comparisons, are presented, with t-values shown in red for positive effects and blue for negative effects.

A nonparametric cluster-based permutation analysis indicated a significant main effect of age group (cluster-t = 7434, p < 0.001), with older adults showing lower mPAC than younger adults. This effect corresponded to a cluster in the observed data spanning all time points and frequencies from 30 to 80 Hz. Next, the CBPT analysis indicated a significant main effect of repetition (cluster-t = 5648.2, p < 0.001), with mPAC values increasing across sequence repetitions. The effect corresponded to a positive cluster in the observed data spanning all time points and frequencies from 30 to 100 Hz. Moreover, the CBPT analysis indicated a significant main effect of performance (cluster-t = 9582.9, p < 0.001), with lower mPAC observed in slower learners compared to faster learners. The effect corresponded to a negative cluster spanning all time points and frequencies from 30 to 100 Hz. Furthermore, the CBPT analysis indicated a significant interaction effect between age group and performance (cluster-t = 2901.3, p = 0.003), with younger adults showing a stronger association between higher mPAC and better performance. This effect corresponded to a positive cluster spanning all time points and frequencies from 30 to 90 Hz. The CBPT analysis also indicated a significant interaction effect between repetition and performance (cluster-t = 359.9, p = 0.031), with faster learners showing more substantial increases in mPAC with repetition. This effect corresponded to a negative cluster in the observed data around 0.3 to 0.9 seconds, within the frequency ranges of 40 to 60 Hz and 80 to 100 Hz. Finally, the CBPT analysis indicated a significant three-way interaction effect of age group, repetition, and performance (cluster-t = 994.8, p = 0.011). This effect corresponded to a positive cluster in the observed data around 0 to 0.9 seconds and frequencies from 40 to 90 Hz, indicating that the combined influence of age, repetition, and learning speed significantly impacts mPAC, with younger, faster learners showing the strongest increases of mPAC.

Next, to visualize the effects obtained from the CBPT procedure, we selected a time window from 0 to 0.9 seconds and a frequency range from 30 to 80 Hz, where the primary effects of interest were consistently observed. We then averaged the mPAC values across these selected time points and frequencies and plotted them as an error bar plot across sequence repetitions and performance groups (Figure 4 C & D). The plot shows that mPAC values vary across age groups, repetition number, and performance groups. Specifically, older adults have lower mPAC values than younger adults across repetitions, and mPAC generally increases with repetition, reflecting the main effects observed in the CBPT analysis. Additionally, faster learners exhibit higher mPAC compared to slower learners, with the interaction effects indicating that younger, faster learners show the most substantial increases in mPAC with repetition.

**Figure 4:**
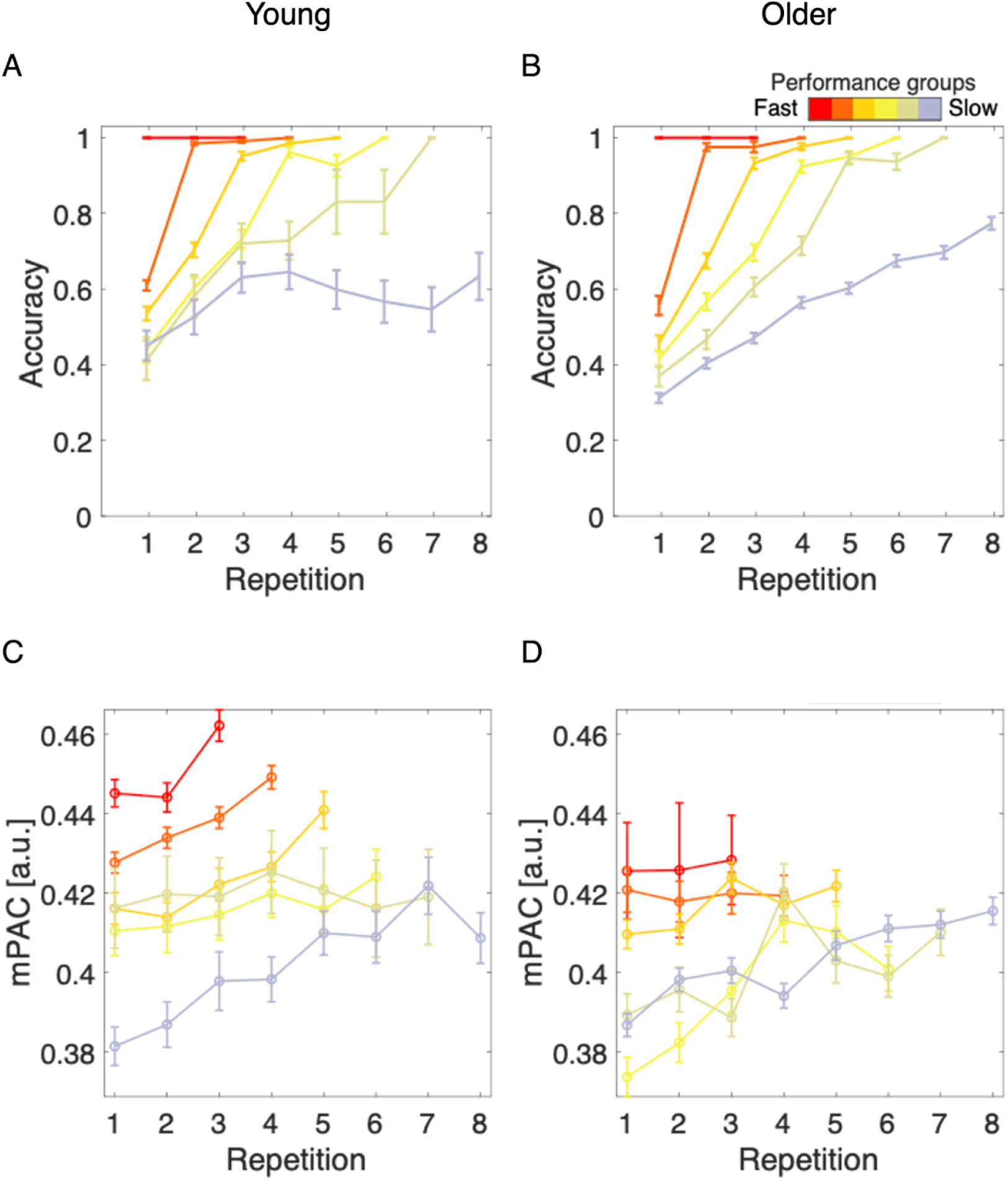
Behavioral performance in the sequence learning task (top panel) and multivariate phase-amplitude coupling (mPAC) over sequence repetitions in young (A & C) and older (B & D) participants. Panel A & B: The task lasted 3 to 8 repetitions, depending on learning rate. The lines indicate the learning curves of participants that completed the task after 3, 4, 5, 6, 7 or 8 sequence repetitions. Panel C & D: The mPAC was computed for a time window of 0 to 900 ms post stimulus and frequency range 30 to 80 Hz. The lines represent mPAC progression of subjects that completed the task after 3 to 8 sequence repetitions (performance groups). Error bars represent the standard error of the mean.

To ensure that the observed mPAC effects were not simply due to variations in the number of stimuli across sequences (i.e., fast learners completing only 3 repetitions, while slow learners completed up to 8), we performed a set of supplementary sensitivity analyses, varying the number of stimuli used to compute GED (Supplementary Material 4). In the first approach, we limited the PAC computation to the stimuli from the first three repetitions (24 stimuli), as all subjects completed at least three sequence repetitions. Using these stimuli, we applied GED to extract the mid-frontal theta component with the highest eigenvalue. This theta component was then applied across all repetitions within each sequence. The mid-frontal theta troughs from all these repetitions were subsequently used to perform a second GED, from which the theta trough component was extracted. In an alternative approach, we computed both the initial GED for the mid-frontal theta component and the second GED for the theta trough component using only the first three repetitions (24 stimuli). While this analysis minimized potential variation due to unequal stimulus counts, it also reduced power due to the limited number of stimuli. To improve power slightly, we conducted an additional analysis using the first four repetitions (32 stimuli) for both GED computations, as most subjects completed at least four repetitions. This approach also enhanced the signal-to-noise ratio, as additional stimuli improved the stability of the PAC estimates. Importantly, we applied this process consistently across both stages of GED computation - first to isolate the mid-frontal theta component and then to extract the mid-frontal theta trough component (Supplementary Material 4).

Critically, across all these control analyses, we consistently observed the same main effects of age group, repetition, and performance and the interaction of age group and performance. In analyses with a reduced number of stimuli, the interaction between repetition and performance, as well as the three-way interaction of age group, repetition, and performance did not reach statistical significance. Using fewer stimuli reduced statistical power, making it more challenging to detect finer interaction effects, particularly those involving repetition and performance. This expected reduction in power reflects the inherent trade-off of using fewer stimuli, resulting in marginally attenuated effects in the subset analyses. Nevertheless, these control analyses support the robustness of our findings, showing that the primary effects are not simply artifacts of the number of stimuli included in each participant’s sequence.

### 3.3. Multivariate mid-frontal theta and gamma over the course of learning and their relation to mPAC

One important consideration is whether the mPAC effects might simply reflect variations in mid-frontal theta and gamma power over the course of learning. To explore this possibility, we examined mid-frontal theta and gamma power, derived from the mid-frontal theta and gamma components identified through GED, over the course of learning. Specifically, mid-frontal theta and gamma power were averaged over the same time window as mPAC (0 to 0.9 seconds) to ensure comparability. We then computed a linear mixed-effect model to test the effects of age group, repetition, performance, as well as all interaction on multivariate mid-frontal theta.

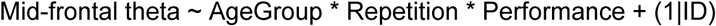

The model revealed a significant main effect of age group (β =- 0.10; p = 7.3e-14; CI = [−0.12, −0.07]), suggesting a lower mid-frontal theta in the older group. Moreover, there was a significant main effect of repetition number, (β = 0.01; p = 6.7e-4; CI = [0.01, 0.02]) that is the mid-frontal theta increased with each sequence repetition. There was also a significant main effect of performance (β = −0.04; p = 6.8e-16; CI = [−0.05, −0.03]), indicating lower mid-frontal theta in slow learners. Additionally, there was a significant interaction of age group and performance (β = 0.02; p = 0.01; CI = [0.01, 0.03]), indicating that the difference between fast and slow learners was smaller in older compared to young subjects. The interactions between age group and repetition number (β = −0.00; p = 0.93; CI = [−0.01, 0.01]), repetition number and performance (β = 0.00; p = 0.38; CI = [−0.01, 0.01]), as well as the three-way interaction of age group, repetition number and performance (β = 0.00; p = 0.77; CI = [−0.01, 0.01]) did not reach statistical significance. In addition, we observed a substantial variation of accuracy between subjects (SD = 0.09).

Next, we computed a linear mixed-effect model to test the effects of age group, repetition, performance, as well as all interaction on gamma.

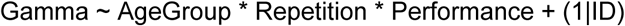

The model revealed a significant main effect of age group (β = 0.01; p = 0.044; CI = [0.00, 0.02]), suggesting a higher gamma in the older group. Moreover, there was a significant main effect of repetition number, (β = 0.01; p = 0.042; CI = [0.00, 0.02]) that is the gamma increased with each sequence repetition. There was also a significant main effect of performance (β = −0.02; p = 8.7e-5; CI = [−0.02, −0.01]), indicating lower gamma in slow learners. The interactions between of age group and performance (β = 0.01; p = 0.255; CI = [−0.01, 0.02]), age group and repetition number (β = −0.01; p = 0.254; CI = [−0.02, 0.01]), repetition number and performance (β = 0.00; p = 0.698; CI = [−0.01, 0.01]), as well as the three-way interaction of age group, repetition number and performance (β = 0.00; p = 0.50; CI = [−0.01, 0.02]) did not reach statistical significance.

Finally, we examined whether differences in mPAC were still evident across performance groups when mid-frontal theta and gamma power were matched (see Methods 2. 6. 3). The stratification analysis was conducted separately for young and older adults, focusing on the three performance groups with the largest sample sizes for each age group. For young adults, performance groups 3, 4, and 5 were analyzed, while for older adults, performance groups 5, 6, and 8 were included. Our findings revealed that even among subjects with similar mid-frontal theta power and similar gamma power, differences in mPAC persisted across performance groups. Specifically, high performing subjects continued to exhibit stronger mPAC compared to low performing subjects, despite comparable mid-frontal theta or gamma power levels. These results highlight the robustness of the mPAC effect as being independent of mid-frontal theta and gamma power (see Figure 5).

**Figure 5:**
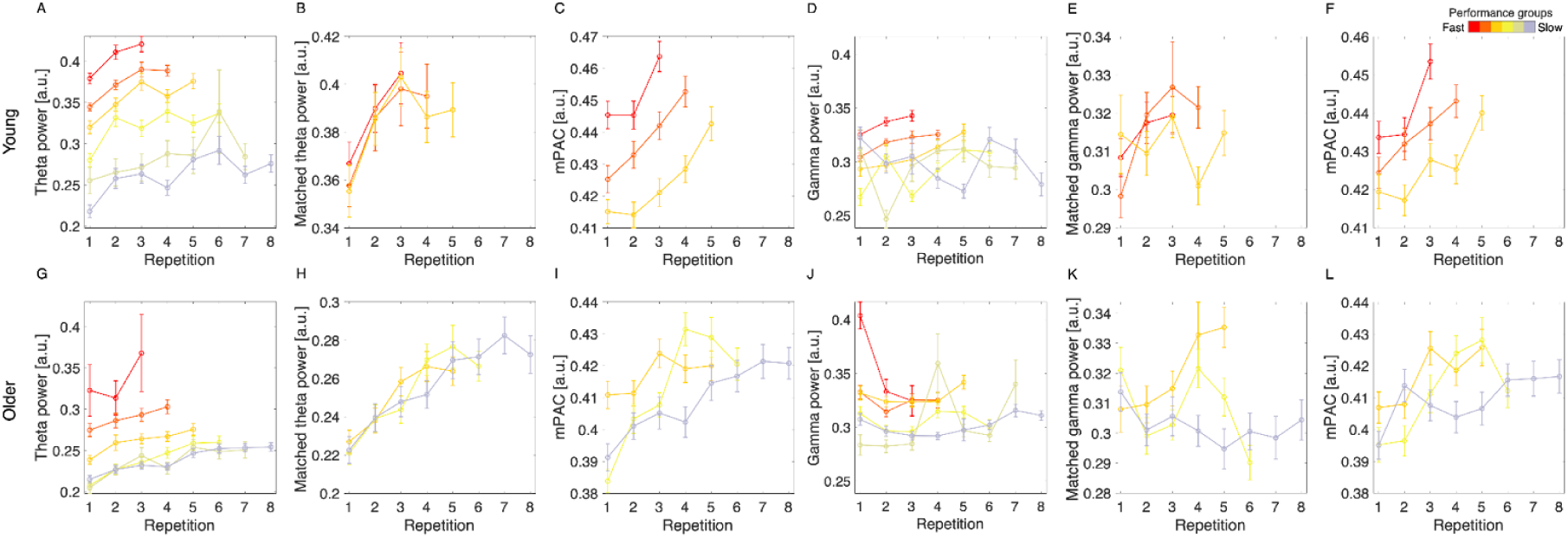
Multivariate mid-frontal theta (A & G) and gamma (D & J) over sequence repetitions and the results of stratification analyses in young (top panel) and older (bottom panel) subjects. (B & H) Matched mid-frontal theta power across three performance groups. As increasing the number of performance groups for this analysis reduces the pool of subjects with matching mid-frontal theta power levels, we focused on only the three performance groups with the largest sample sizes. (C & I) mPAC over sequence repetitions for subjects with matched mid-frontal theta levels. (E & K) Matched gamma power across three performance groups. (F & L) mPAC over sequence repetitions with matched gamma levels. Error bars represent the standard error of the mean.

## 4. Discussion

In this study, we explored age-related differences in memory formation processes using multivariate mid-frontal theta-gamma phase-amplitude coupling through generalized eigendecomposition. With this approach, we addressed the limitations of traditional univariate methods, including bias estimates from non-sinusoidal oscillations and a focus on individual electrodes, enabling a broader assessment of theta-gamma coupling in coordinating activity across brain regions. Utilizing a visual sequence learning paradigm in which participants learned a fixed sequence of visual stimuli over repeated observations, we found that younger participants learned faster compared to older participants, highlighting age-related declines in processing speed and working memory efficiency. Neurophysiological data revealed that mPAC increased progressively over the course of the learning process in both age groups. However, older adults demonstrated lower mPAC strength overall, as did slow learners within both age groups. Critically, stratification analysis indicated that mPAC effects persisted across performance groups with similar mid-frontal theta activity, emphasizing that mPAC provides unique insights beyond theta contributions alone. These findings advance our understanding of the neural underpinnings of age-related memory decline and present avenues for targeted interventions to support memory performance.

### Multivariate theta-gamma phase-amplitude coupling over the course of learning

The theoretical model proposed by Lisman and Idiart (1995) suggests that individual items to be maintained in WM are represented by single gamma cycles. These gamma cycles are nested within a theta cycle, creating a hierarchical framework for neural coding. Theta-gamma coupling enables the sequential organization of these items, allowing the brain to effectively manage, integrate, and bind them into cohesive memory traces (Jensen & Lisman, 1996; J. Lisman & Jensen, 2013; Sauseng et al., 2009, 2019). Since only a finite number of gamma cycles can fit within a single theta cycle, this mechanism provides a plausible explanation for the limited capacity of working memory (Cowan, 2001; Jensen & Lisman, 1996; Sauseng et al., 2019).

Our study revealed that younger subjects exhibit higher overall mPAC strength compared to older subjects, indicative of better neural coordination between mid-frontal theta and occipital gamma oscillations during our visual learning task. This finding aligns with previous research suggesting that aging is associated with reduced neural plasticity, affecting learning and memory processes, including visual learning (Antonenko & Flöel, 2014; Werkle-Bergner et al., 2009). Moreover, Patterson et al. (2015) showed that decreased theta-gamma coupling is related to decreased performance in older age. The diminished mPAC strength in older adults may reflect less efficient neural coordination during visual learning, contributing to their decreased performance (Antonenko & Flöel, 2014; Karlsson et al., 2022; Patterson et al., 2024; Sweeney-Reed et al., 2015; Werkle-Bergner et al., 2009). In our supplementary analyses (Supplementary Material 1. 8), we demonstrated that older subjects exhibit significantly faster theta rhythms compared to young subjects. This faster theta limits the number of gamma cycles that can fit within a single theta cycle. This observation aligns with findings from experimental modulation of theta frequency using transcranial alternating current stimulation, which further supports this relationship (Vosskuhl et al., 2015; Wolinski et al., 2018). Specifically, slowing theta rhythms enhances working memory capacity by accommodating more gamma cycles, while faster theta rhythms diminish capacity by limiting gamma nesting.

Fast learners in our study exhibited higher overall mPAC strength than slow learners, indicating that individuals who perform better in visual learning tasks tend to have stronger phase-amplitude coupling, reflecting more efficient neural processing. Stronger PAC in fast learners aligns with findings from previous research. Numerous studies have demonstrated that enhanced theta-gamma coupling is associated with improved learning of item-context associations and higher working memory capacity (Chaieb et al., 2015; Karlsson et al., 2022; Sauseng et al., 2009, 2019; Staudigl & Hanslmayr, 2013; Tort et al., 2009). Specifically, Chaieb et al. (2015) showed that increased theta-gamma PAC correlates with the ability to maintain multiple items in WM, highlighting its role in supporting memory capacity. Similarly, Tort et al. (2009) demonstrated that theta-gamma coupling strengthens during learning tasks, exhibiting a direct relationship with memory performance through the encoding of item-context associations. Further evidence supporting the role of PAC in cognitive enhancement comes from studies employing non-invasive brain stimulation (Diedrich et al., 2024; Reinhart & Nguyen, 2019). For instance, Reinhart and Nguyen (2019) conducted a randomized, sham-controlled, double-blind study in which participants received transcranial alternating current stimulation targeting theta and gamma oscillations while performing working memory tasks. The results revealed increased accuracy and improved WM performance, indicating that stimulating and synchronizing neural oscillations can enhance cognitive functions related to visual learning and memory.

Our findings also showed that mPAC values increased significantly with sequence repetitions. Repetitive visual training strengthens neural connections and enhances synchronization within the visual cortex, facilitating more efficient processing of visual information (Voytek et al., 2010). Notably, mPAC increased over the course of learning, with the most pronounced changes observed in young fast learners. This highlights how repeated engagement enhances oscillatory synchronization most effectively in individuals with higher learning capacity, emphasizing the role of neural plasticity and effective practice in optimizing visual learning.

Importantly, our results suggest that young subjects and fast learners exhibit elevated mPAC strength that extends beyond event-related (post-stimulus) activity, persisting throughout the entire epoch, including the pre-stimulus interval. This finding indicates that successful learning is less reliant on event-specific neural activity and more dependent on overall cognitive preparedness or memory capacity, reflected in higher baseline mPAC levels. Furthermore, in previous work (Strzelczyk & Langer, 2024), we have shown that pre-stimulus mid-frontal theta activity is closely linked to memory and learning and is influenced by aging, with theta power increasing over the course of learning in both younger and older individuals but at a sharper rate in younger participants. This highlights age-related differences in the ability to enter preparatory neural states essential for effective memory encoding and retrieval. Taken together, these findings highlight the critical role of baseline neural activity in learning and memory formation, supporting the idea that resting-state brain dynamics can predict learning performance.

In the present study, we employed multivariate phase-amplitude coupling analysis via generalized eigendecomposition to address several limitations of traditional univariate PAC methods (Cohen, 2017b). This approach is particularly robust against the non-stationarity of EEG signals, reducing spurious or biased estimates and enhancing the signal-to-noise ratio (SNR) by integrating data across multiple electrodes (Cohen, 2017b, 2022). This improvement is especially critical for detecting signals with inherently low amplitudes, such as high-frequency gamma power, which are not only weak but also prone to contamination by noise and artifacts. By incorporating a spatial filter that computes weighted averages from multiple electrodes, the multivariate method isolates specific patterns of activity that would otherwise remain obscured in single-electrode analyses due to noise contamination. Univariate PAC methods typically analyze coupling within individual electrodes, overlooking the broader functional role of theta-gamma interactions in coordinating activity across distributed brain regions. This limitation is particularly significant given the distinct origins of theta and gamma oscillations - theta oscillations primarily originate in the hippocampus, while gamma oscillations are often associated with occipital cortex activity during tasks involving visual processing (Axmacher, Cohen, et al., 2010; Axmacher, Henseler, et al., 2010; Buzsaki, 2011; Buzsáki & Wang, 2012; Karlsson et al., 2022; Sauseng et al., 2019). Consequently, traditional methods fail to fully capture the spatially distributed dynamics critical for understanding theta-gamma interactions in memory and learning processes (Papaioannou et al., 2022; Sauseng et al., 2009).

The multivariate approach is distinct in its ability to identify a spatial filter that maximizes the contrast between theta troughs and broadband. Instead of examining PAC at predefined electrodes, we derived a data-driven theta trough component for each subject, representing the neural patterns most strongly associated with mid-frontal theta troughs. Importantly, we did not impose constraints on the brain region or frequency band, allowing the analysis to adaptively identify the spatial filter that maximizes this contrast. This fully data-driven approach revealed that the maximal difference between theta trough and other phases was localized in occipital brain regions, aligning with gamma bursts expected during visual item processing. By accounting for inter-individual differences in theta topography, the multivariate approach enables more precise isolation of subject-specific neural patterns. The occipital topography of the theta trough component observed across participants further highlights the relevance of this approach in linking theta-gamma coupling to visual and memory-related processes.

### Multivariate mid-frontal theta and gamma power over the course of learning

So far we established that mPAC effects change significantly over the course of learning, but these changes might merely reflect variations in mid-frontal theta or gamma power. To explore this possibility, we assessed multivariate mid-frontal theta and gamma power dynamics over the course of learning, leveraging the components identified via GED. Our analyses revealed that mid-frontal theta power patterns closely resemble those of mPAC during learning, exhibiting similar significant effects. Critically, sensitivity analyses demonstrated that mPAC effects are not solely driven by mid-frontal theta power increases. When mid-frontal theta power was included as a covariate in the model, the effects of age group, repetition and performance, as well all interactions remained significant, indicating that mPAC captures variance beyond mid-frontal theta power (Supplementary Material 1. 6). Stratification analyses further supported this finding, showing that mPAC can distinguish fast and slow learners with similar mid-frontal theta and gamma power levels, highlighting its unique contribution to memory processes.

Mid-frontal theta has long been associated with memory formation, with studies reporting increased activity during the encoding of successfully remembered items compared to those forgotten (Herweg et al., 2020; Klimesch et al., 2001; Sauseng et al., 2009; Schreiner & Staudigl, 2020; Staudigl et al., 2010; Staudigl & Hanslmayr, 2013; Strzelczyk & Langer, 2024). Additionally, individuals with higher cognitive performance tend to maintain higher gamma power in parieto-occipital areas, suggesting that preserved posterior-occipital gamma activity is associated with better cognitive function (Bakhtiari et al., 2023). Moreover, older adults exhibit reduced gamma activity, and age-related reductions in gamma oscillations correlate with declines in visual short-term memory (Werkle-Bergner et al., 2009). Yet, some argue that while mid-frontal theta is necessary for memory formation, it is not sufficient. Studies have found no significant age-related differences in theta power between younger and older adults, whereas PAC exhibited clear age effects (Karlsson & Sander, 2023). This suggests that memory processes may not be driven solely by theta or gamma oscillations, but rather by the precise coordination of occipital gamma activity with mid-frontal theta power. Previously, we showed that univariate mid-frontal theta power was reduced in older adults compared to younger adults (Strzelczyk & Langer, 2024). However, univariate mid-frontal theta power alone was insufficient to differentiate between fast and slow learners (Supplementary Material 1. 7). Only by employing a multivariate approach, which utilizes data from all electrodes to enhance SNR, were we able to discriminate between fast and slow learners effectively. Our findings suggest that mid-frontal theta power reductions in older adults and slow learners reflect diminished neural efficiency in coordinating memory-related processes. However, these changes do not fully account for individual differences in learning and memory. Critically, the coordination between mid-frontal theta and occipital gamma activity emerges as a central mechanism driving successful memory formation. Our results support the idea that both mid-frontal theta power and cross-frequency coupling are essential for successful learning, indicating that the combination of theta oscillations and their coordination with occipital gamma activity better explains individual differences in learning performance and age-related changes in memory.

## Conclusion

In conclusion, our study provides novel insights into the neurophysiological mechanisms contributing to successful learning across young and older participants. Our data showed that young subjects exhibit higher mPAC strength compared to older subjects. Importantly, mPAC successfully distinguished between fast and slow learners across both age groups, highlighting its role in memory formation processes. Crucially, our stratification analysis demonstrated that mPAC effects persisted even when controlling for mid-frontal theta and gamma power, indicating that mPAC is not merely a byproduct of theta and gamma amplitude but instead reflects a distinct mechanism supporting memory formation. Taken together, these findings shed light on age-related differences in memory formation and emphasize the role of mPAC in learning efficiency. By identifying neural markers that differentiate fast from slow learners, this work may inform targeted interventions to enhance memory performance in older adults and individuals with learning difficulties. Future research could explore whether modulating mPAC through neurostimulation techniques, such as TMS could improve learning outcomes, particularly in populations with impaired cognitive function.

