## Supplement for "Impact of Aging on Theta-Phase Gamma-Amplitude Coupling During Learning: A Multivariate Analysis"

### 1. Supplementary Material

#### 1. 1. Reaction times over the course of learning

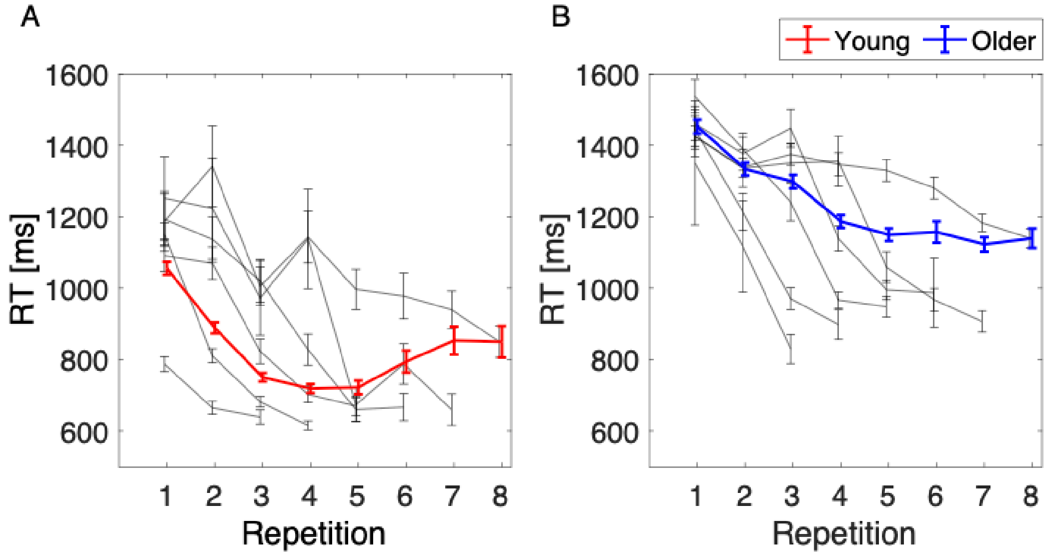


Supplementary Figure 1: Reaction times in young (A) and older (B) participants. Please note that the participants were not explicitly instructed to perform the task as fast as possible. Hence, the reaction times need to be interpreted with caution.

1. 2. Multivariate theta-gamma phase-amplitude coupling with filter of 5Hz instead of 10 Hz

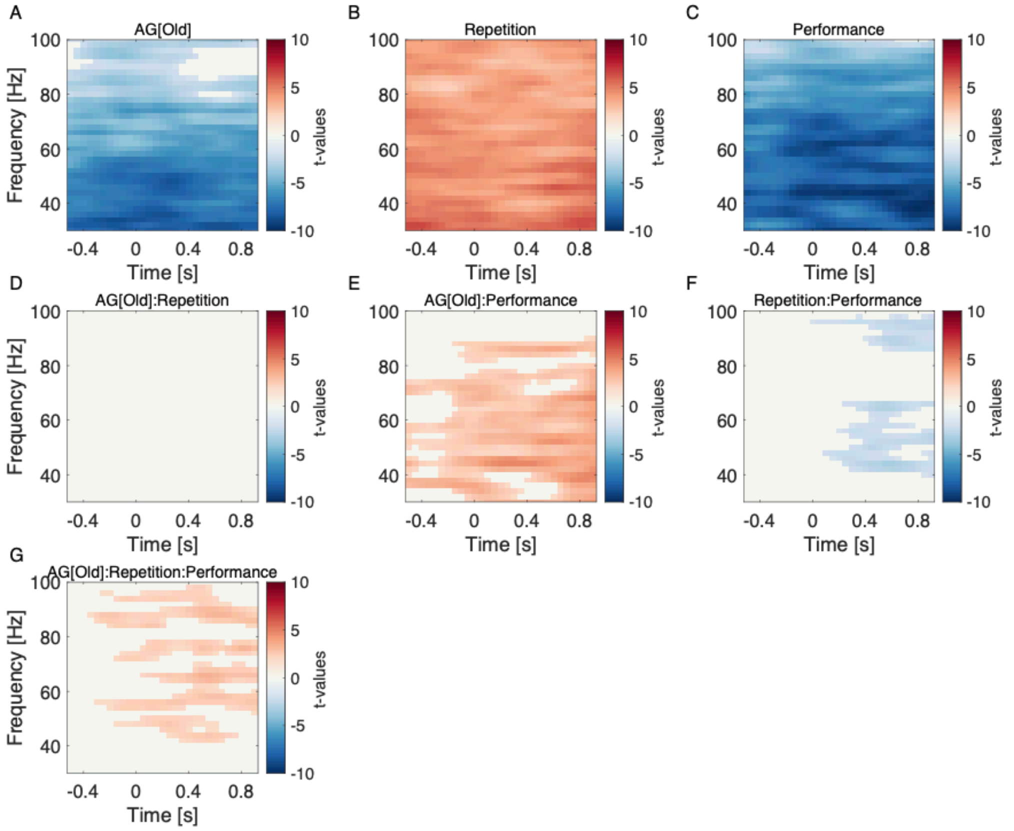


Supplementary Figure 2: Results of mass linear mixed-effects modeling of multivariate phase-amplitude coupling with crossed random effects. Cluster-based permutation analysis indicated that (A) older adults exhibited lower mPAC compared to younger adults, (B) mPAC values increased across sequence repetitions, (C) slower learners showed lower mPAC compared to faster learners, (E) a significant interaction between age group and performance indicated a stronger relationship between higher mPAC and better performance in younger adults, (F) faster learners demonstrated greater increases in mPAC with repetition, (G) a three-way interaction of age group, repetition, and performance revealed that younger, faster learners had the most pronounced increases in mPAC. Only statistically significant clusters, corrected for multiple comparisons, are presented, with t-values shown in red for positive effects and blue for negative effects.

1. 3. Effect of sliding window length on multivariate theta-gamma phase-amplitude coupling

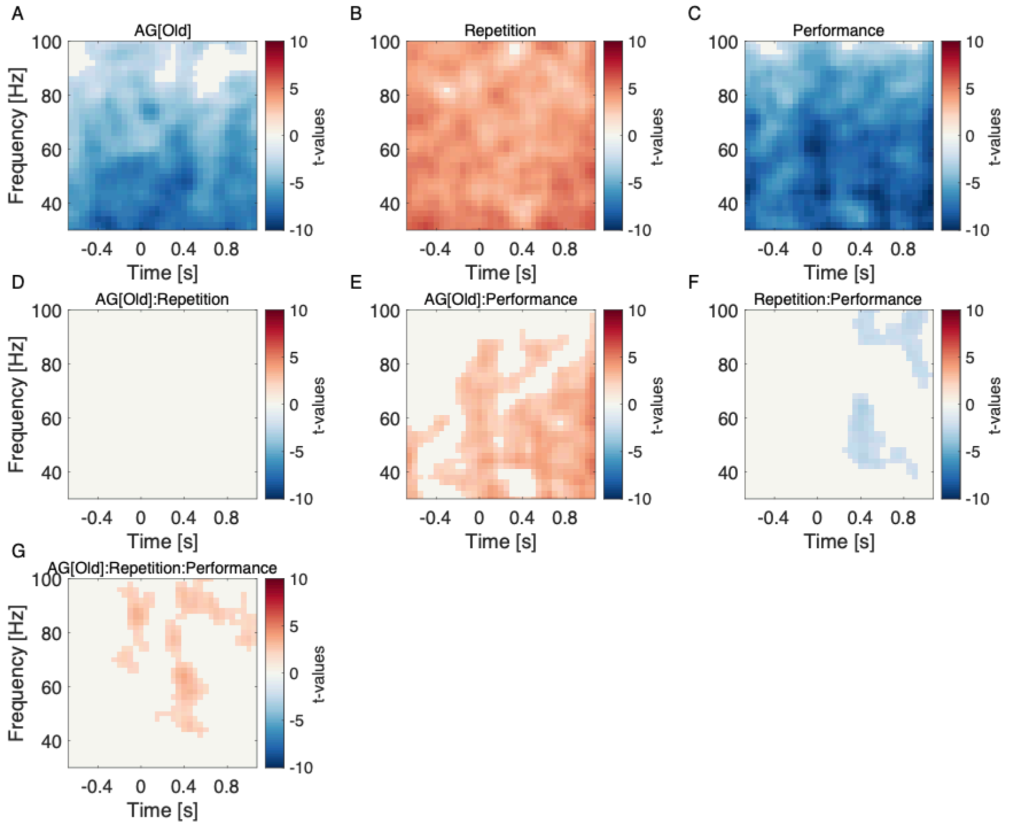


Supplementary Figure 3: Results of mass linear mixed-effects modeling of multivariate phase-amplitude coupling with crossed random effects, but with a sliding window of 200 ms instead of 500 ms. Cluster-based permutation analysis indicated that (A) older adults exhibited lower mPAC compared to younger adults, (B) mPAC values increased across sequence repetitions, (C) slower learners showed lower mPAC compared to faster learners, (E) a significant interaction between age group and performance indicated a stronger relationship between higher mPAC and better performance in younger adults, (F) faster learners demonstrated greater increases in mPAC with repetition, (G) a three-way interaction of age group, repetition, and performance revealed that younger, faster learners had the most pronounced increases in mPAC. Only statistically significant clusters, corrected for multiple comparisons, are presented, with t-values shown in red for positive effects and blue for negative effects.

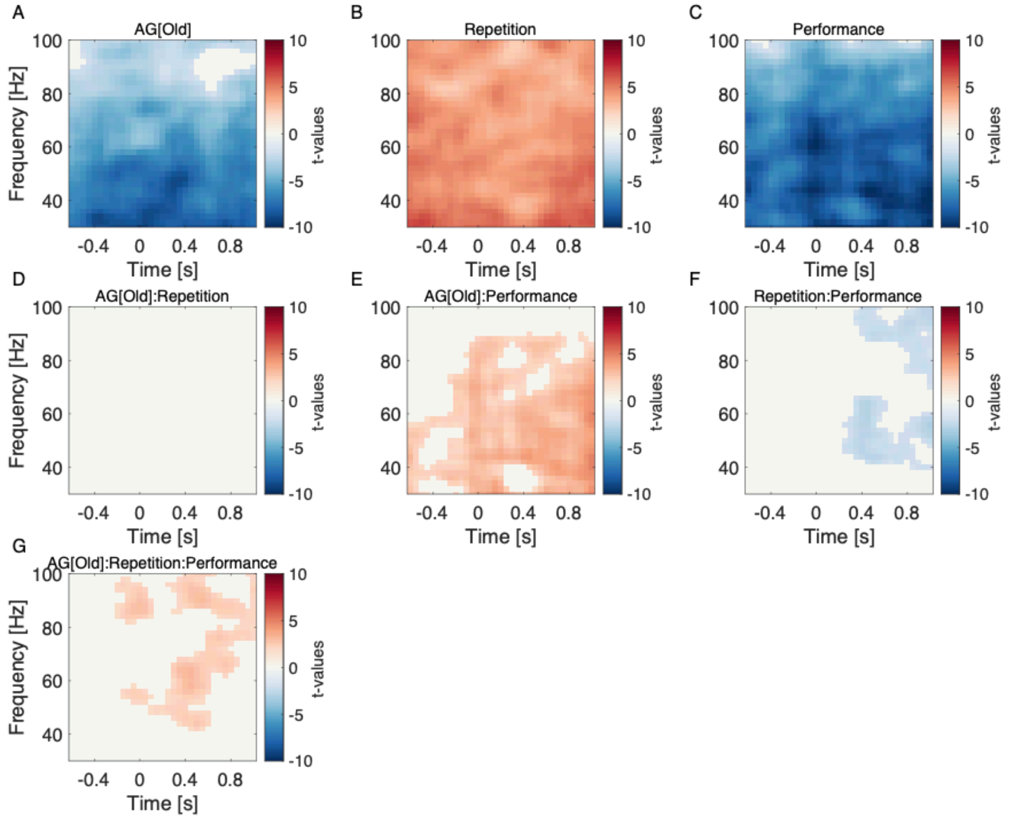


Supplementary Figure 4: Results of mass linear mixed-effects modeling of multivariate phase-amplitude coupling with crossed random effects, but with a sliding window of 300 ms instead of 500 ms. Cluster-based permutation analysis indicated that (A) older adults exhibited lower mPAC compared to younger adults, (B) mPAC values increased across sequence repetitions, (C) slower learners showed lower mPAC compared to faster learners, (E) a significant interaction between age group and performance indicated a stronger relationship between higher mPAC and better performance in younger adults, (F) faster learners demonstrated greater increases in mPAC with repetition, (G) a three-way interaction of age group, repetition, and performance revealed that younger, faster learners had the most pronounced increases in mPAC. Only statistically significant clusters, corrected for multiple comparisons, are presented, with t-values shown in red for positive effects and blue for negative effects.

1. 4. Effect of trial number on multivariate theta-gamma phase-amplitude coupling

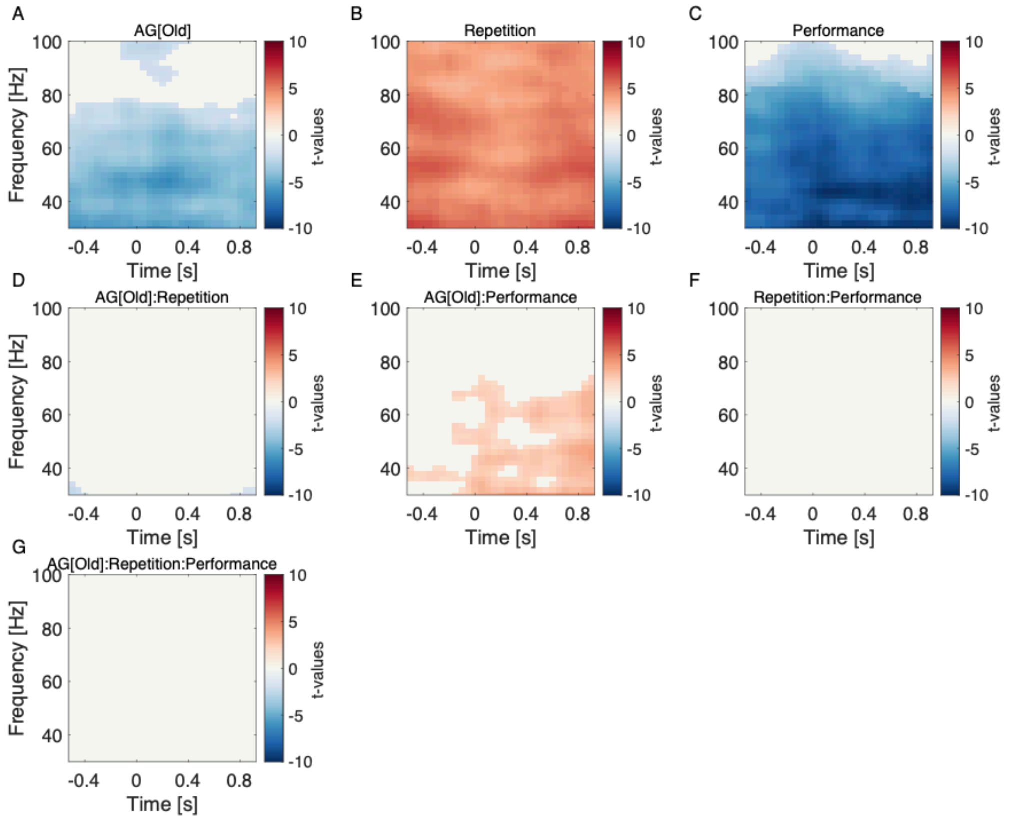


Supplementary Figure 5: Results of mass linear mixed-effects modeling of multivariate phase-amplitude coupling with crossed random effects. First 24 trials from the first 3 sequence repetitions were used in GED to extract the mid-frontal theta component. All trials from all repetitions were used in GED to extract the theta trough component. Cluster-based permutation analysis indicated that (A) older adults exhibited lower mPAC compared to younger adults, (B) mPAC values increased across sequence repetitions, (C) slower learners showed lower mPAC compared to faster learners, (E) a significant interaction between age group and performance indicated a stronger relationship between higher mPAC and better performance in younger adults. Only statistically significant clusters, corrected for multiple comparisons, are presented, with t-values shown in red for positive effects and blue for negative effects.

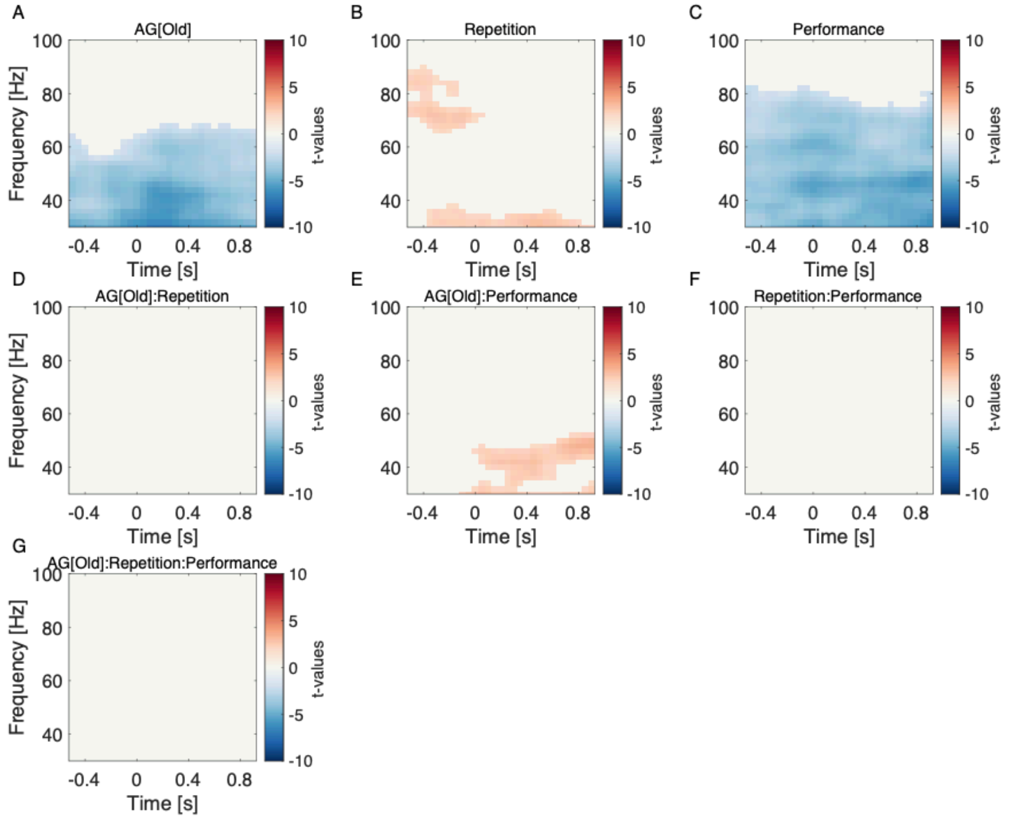


Supplementary Figure 6: Results of mass linear mixed-effects modeling of multivariate phase-amplitude coupling with crossed random effects. First 24 trials from the first 3 sequence repetitions were used in GED to extract the mid-frontal theta component and the theta trough component. Cluster-based permutation analysis indicated that (A) older adults exhibited lower mPAC compared to younger adults, (B) mPAC values increased across sequence repetitions, (C) slower learners showed lower mPAC compared to faster learners, (E) a significant interaction between age group and performance indicated a stronger relationship between higher mPAC and better performance in younger adults. Only statistically significant clusters, corrected for multiple comparisons, are presented, with t-values shown in red for positive effects and blue for negative effects.

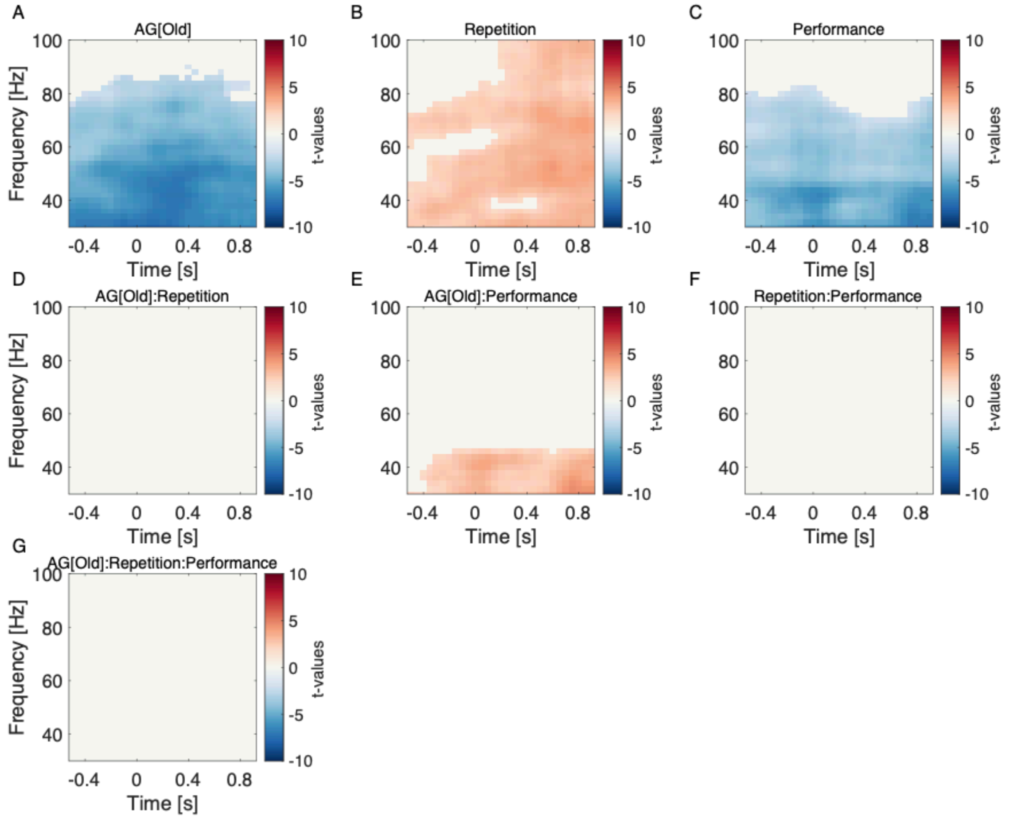


Supplementary Figure 7: Results of mass linear mixed-effects modeling of multivariate phase-amplitude coupling with crossed random effects. First 32 trials from the first 4 sequence repetitions were used in GED to extract the mid-frontal theta component and the theta trough component. Increasing the number of stimuli used in GED increases statistical power to detect an effect. Cluster-based permutation analysis indicated that (A) older adults exhibited lower mPAC compared to younger adults, (B) mPAC values increased across sequence repetitions, (C) slower learners showed lower mPAC compared to faster learners, (E) a significant interaction between age group and performance indicated a stronger relationship between higher mPAC and better performance in younger adults. Only statistically significant clusters, corrected for multiple comparisons, are presented, with t-values shown in red for positive effects and blue for negative effects.

#### 1. 5. Multivariate theta-gamma phase-amplitude coupling with simulated theta-trough component

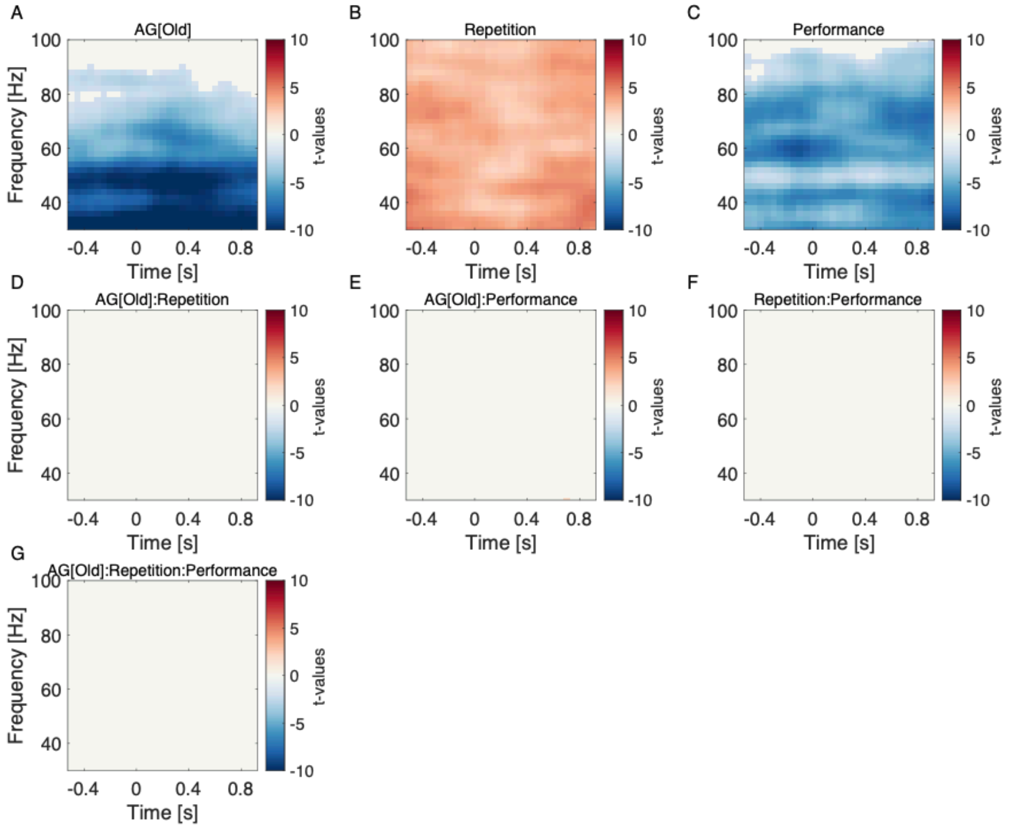


Supplementary Figure 8: Results of mass linear mixed-effects modeling of multivariate phase-amplitude coupling with crossed random effects. The expected occipital gamma was simulated and a GED component with the highest spatial correlation with simulated gamma was selected for mPAC computation. Cluster-based permutation analysis indicated that (A) older adults exhibited lower mPAC compared to younger adults, (B) mPAC values increased across sequence repetitions, (C) slower learners showed lower mPAC compared to faster learners. Only statistically significant clusters, corrected for multiple comparisons, are presented, with t-values shown in red for positive effects and blue for negative effects.

#### 1. 6. Results of mass linear mixed-effects modeling with mid-frontal theta as a covariate

To further illuminate the differences and similarities between mid-frontal theta power and mPAC, we computed the correlation between mid-frontal theta power and mPAC, which yielded a relatively low value (r = 0.17). To account for the nested structure of interactions between repetitions, age groups, performance, and individual subjects, we further conducted mixed model analyses using CBPT to determine whether mid-frontal theta power and mPAC are related when controlling for key variables: repetition number, age group, performance, and a random effect of subject.

$$mPAC_{[\text{time point, frequency}]} \sim AG * Repetition * Performance + \text{Mid-frontal theta} + (1|ID)$$

The results from these analyses indicate that, while mid-frontal theta power is significantly related to mPAC, the main effects of age group, repetition, and performance remain significant. The outcomes of the CBPT procedure are presented in Supplementary Figure 9.

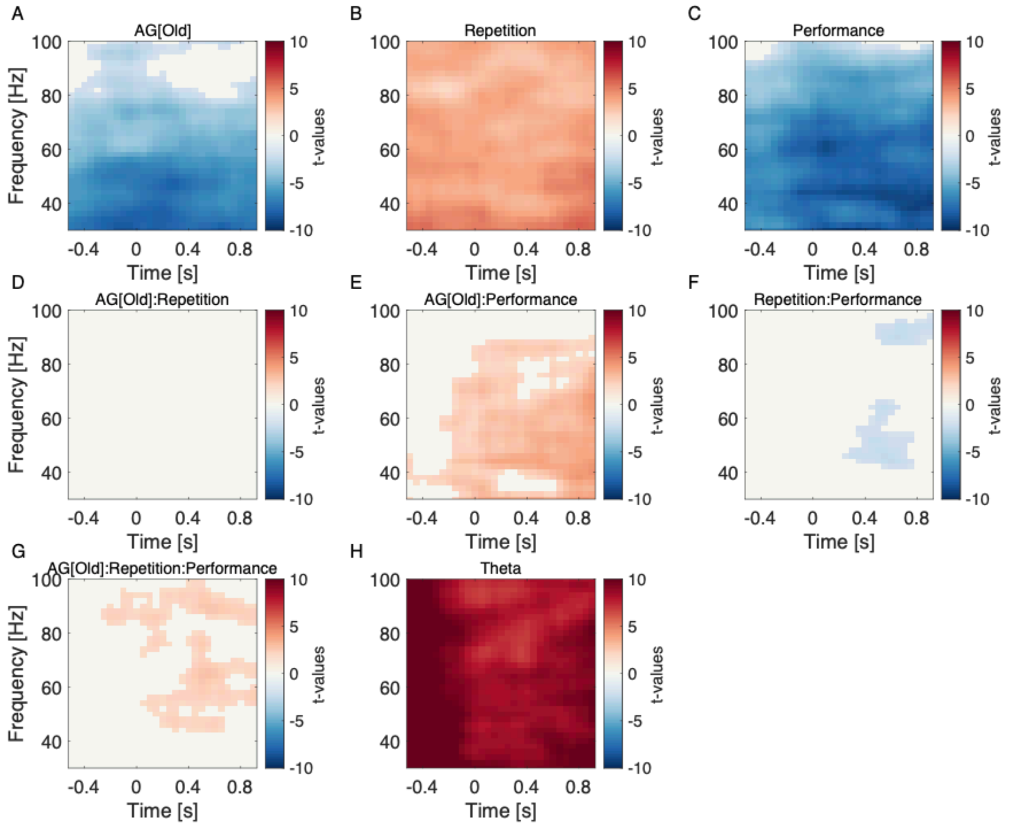


Supplementary Figure 9: Results of mass linear mixed-effects modeling of multivariate phase-amplitude coupling with crossed random effects, but with mid-frontal theta power as a covariate. Cluster-based permutation analysis indicated that (A) older adults exhibited lower mPAC compared to younger adults, (B) mPAC values increased across sequence repetitions, (C) slower learners showed lower mPAC compared to faster learners, (E) a significant interaction between age group and performance indicated a stronger relationship between higher mPAC and better performance in younger adults, (F) faster learners demonstrated greater increases in mPAC with repetition, (G) a three-way interaction of age group, repetition, and performance revealed that younger, faster learners had the most pronounced increases in mPAC. Only statistically significant clusters, corrected for multiple comparisons, are presented, with t-values shown in red for positive effects and blue for negative effects.

1. 7. Univariate theta power over the course of learning

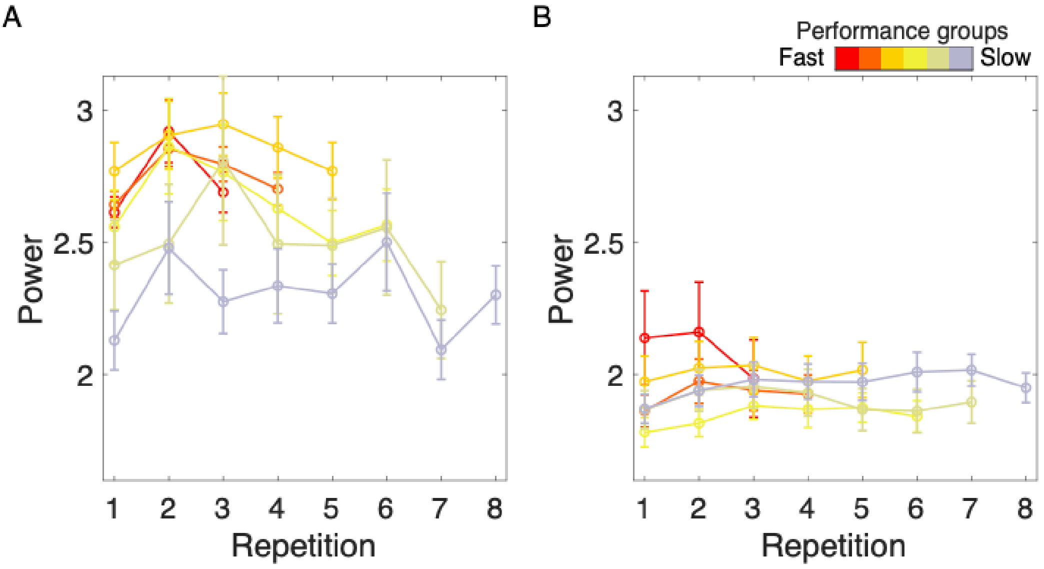


Supplementary Figure 10: Univariate theta power computed using FFT on 6 fronto-central electrodes.

1. 8. Mid-frontal theta peak with highest correlation to simulated mid-frontal theta

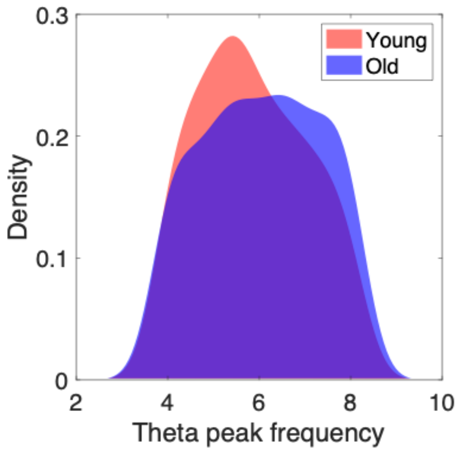


Supplementary Figure 11: Distribution of mid-frontal theta peaks with highest spatial correlation with simulated mid-frontal theta in young and older subjects. Young have on average slower theta frequency than older subjects ($M_{Young}$ = 5.87, $M_{Older}$ = 6.10, t-value = -3.148, df = 1196, p = 0.0017, 95%CI = [-0.37, -0.09]). No differences were found between different performance groups.
